# Multimodal Study of the Effects of Varying Task Load Utilizing EEG, GSR and Eye-Tracking

**DOI:** 10.1101/798496

**Authors:** Jannis Born, Babu Ram Naidu Ramachandran, Sandra Alejandra Romero Pinto, Stefan Winkler, Rama Ratnam

**Affiliations:** Advanced Digital Sciences Center, Illinois at Singapore Pte Ltd., Singapore; Coordinated Science Laboratory, College of Engineering, University of Illinois at Urbana-Champaign, Illinois, USA; Institute of Neuroinformatics, ETH Zürich and University of Zürich, Zürich, Switzerland; National University of Singapore, Singapore; Biological and Life Sciences Division, School of Arts and Sciences, Ahmedabad University, Ahmedabad, Gujarat, India

**Keywords:** Electroencephalography, EEG, galvanic skin response, GSR, eye-tracking, multi-modal data fusion, task loading, task difficulty, cognitive load

## Abstract

**Objective:** The effect of task load on performance is investigated by simultaneously collecting multi-modal physiological data and participant response data. Periodic response to a questionnaire is also obtained. The goal is to determine combinations of modalities that best serve as predictors of task performance.

**Approach:** A group of participants performed a computer-based visual search task mimicking postal code sorting. A five-digit number had to be assigned to one of six different non-overlapping numeric ranges. Trials were presented in blocks of progressively increasing task difficulty. The participants’ responses were collected simultaneously with 32 channels of electroencephalography (EEG) data, eye-tracking data, and Galvanic Skin Response (GSR) data. The NASA Task-Load-Index self-reporting instrument was administered at discrete time points in the experiment.

**Main results:** Low beta frequency EEG waves (12.5-18 Hz) were more prominent as cognitive task load increased, with most activity in frontal and parietal regions. These were accompanied by more frequent eye blinks and increased pupillary dilation. Blink duration correlated strongly with task performance. Phasic components of the GSR signal were related to cognitive workload, whereas tonic components indicated a more general state of arousal. Subjective data (NASA TLX) as reported by the participants showed an increase in frustration and mental workload. Based on one-way ANOVA, EEG and GSR provided the most reliable correlation to perceived workload level and were the most informative measures (taken together) for performance prediction.

**Significance:** Numerous modalities come into play during task-related activity. Many of these modalities can provide information on task performance when appropriately grouped. This study suggests that while EEG is a good predictor of task performance, additional modalities such as GSR increase the likelihood of more accurate predictions. Further, in controlled laboratory conditions, the most informative or minimum number of modalities can be isolated for monitoring in real work environments.

## 1. Introduction

An objective method for determining the effect of task load on performance is useful particularly when such information is required in real-time as when the load changes quickly (Coyne et al., 2009; Chen et al., 2012; Hancock et al., 2013). However, the effects of task loading are inherently multi-dimensional and go beyond cognitive and mental effort (Young et al., 2015) or physical effort (Borg, 1990). For example demands may be temporal and perceptual, with effects that lead to fatigue, frustration, or boredom (Szalma et al., 2004; Epps, 2018). Measuring task load along these dimensions, particularly when internal mental and physical states are not readily accessible or observable, makes for a challenging problem. Measurements usually fall into three broad categories. Behavioral measurements such as various types of eye movement (de Greef et al., 2009) or gross motor behaviors (Boxtel and Jesserun, 1993), subjective measures including self-reporting scales such as the multi-dimensional SWAT (Reid and Nygren, 1988) and NASA Task Load Index questionnaires (Hart and Staveland, 1988), and objective physiological measurements which capture a signal that potentially scales or correlates with task loading (Chen et al., 2012; Lean and Shan, 2012; Young et al., 2015; Charles and Nixon, 2019). Among the last are measures such as electroencephalography (EEG), electromyography (EMG), electrocardiography (ECG), galvanic skin response (GSR), inertial measurements, and speech (Chen et al., 2012). The advantage of physiological measurements lies in their objectivity and, in recent years, in the low-cost and ease of deployment of body-worn sensors.

Many studies have measured task loading using a limited number of physiological signals such as eye movement, or EEG, or GSR in an attempt to measure task loading along one or few dimensions (Smallwood and Schooler, 2006; Feng et al., 2013; Lean and Shan, 2012; Charles and Nixon, 2019). However, few studies have combined modalities so that estimation error can be reduced while classification accuracy can be increased (Chen et al., 2012). In an early attempt by Kittler et al. (1998) multimodal data fusion helped to disambiguate data from single modalities, resulting in higher precision and greater reliability. Similarly, Lazzeri et al. (2014) reported the advantage of combining behavioral and psycho-physiological responses in a case study involving robots used in affective communication.

The relation between task load and performance has been extensively studied. For example, Kim (2005) focused on the discrepancy between expected (objective) and experienced (subjective) difficulty. In a collective intelligence regime, Wagner and Suh (2013) found that tasks of medium-range difficulty are suited best for expertise transfer and collective judgments. Adler and Benbunan-Fich (2015) investigated the effect of multitasking with varying subjective difficulty on performance. They found that when the primary task was considered difficult, subjects who were forced to multitask had significantly reduced performance, whereas in the case of an easy primary task, the forced subjects even experienced a performance boost compared to non-multitaskers. Horvath et al. (2006) found that task difficulty was positively correlated with the level of interest an individual had in the task and goal orientation. In practical situations, where subjective and/or objective measures of difficulty may be absent, it remains to find neurophysiological quantities with predictive power for task difficulty.

In the following sections, we present results from an experiment where cognitive task load is gradually varied. The task performance and objective measures obtained simultaneously from EEG, GSR, and eye gaze patterns, are linked with subjective measures of perceived levels of task loading reported by the NASA TLX questionnaire. An earlier report details preliminary findings (Ramachandran et al., 2017).

## 2. Methods

All human subject experiments were conducted at the Advanced Digital Sciences Center (Singapore) and approved by the National University of Singapore (IRB NUS B-15-038). Written informed consent was obtained from all participants.

### 2.1. Participants

A cohort of healthy human subjects (6M/2F, 24-55 years old) was selected to participate in the study. Participants engaged in a computer task simulating a numerical postal code sorting task while they were monitored through non-invasive, physiological techniques (i.e., objective measurements). Participants were also asked to respond to an electronic questionnaire at selected intervals (i.e., subjective measurements). All participants had normal or corrected-to-normal vision and participated voluntarily in the experiment.

### 2.2. Measurements

Participant testing was carried out in a large office room with 80 lux illumination and background sound level of about 60 dB SPL. During the task, scalp-based electroencephalography (EEG) data, galvanic skin response (GSR, i.e., electrodermal activity), and eye movements were recorded, along with a video recording of the participant performing the task (see further below for details on the sensors and instrumentation).

A schematic of the experimental setup is shown in Figure 1. EEG data were collected using a 32-channel ASALab system (ANT Neuro) with a 32-channel EEG cap (Waveguard) which utilizes the 5 percent electrode placement system (an extension of the 10/20 and 10/10 systems). Raw EEG data along with GSR data were sampled at 2.5 kHz. Eye movements were monitored for both eyes independently using an eye-tracking system (SMI REDn Scientific eyetracker, SensoMotoric Instruments GmbH) with a sampling rate of 30 Hz, controlled by SMI Experiment Center software. Participant responses were collected using a 7-button response pad (Cedrus RB730, Cedrus Corp). GSR, eye-movement data, and participant response data were synchronized with EEG data capture. A digital webcam (Logitech C920) was used for videography of the participant during the task, but the video data were not analyzed and are not presented here. Stimulus presentation and response registration (through the response pad) was controlled by SuperLab (Cedrus Corp). Participants were also required to complete an electronic questionnaire that implemented the NASA Task Load Index (NASA-TLX) (Hart and Staveland, 1988). The NASA-TLX self-reporting instrument is a set of questions targeted at mental workload, physical workload, temporal workload, performance, effort, and frustration. The ratings provided by the subjects are subjective, and used to compare and correlate with the objective physiological measurements.

**Figure 1:**
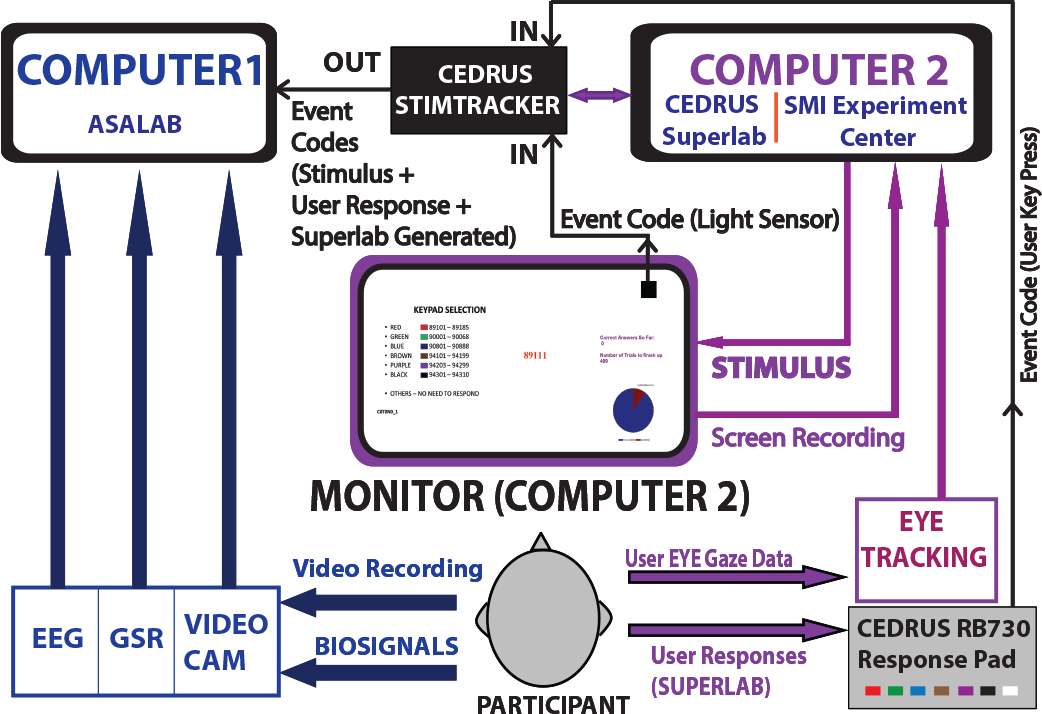
Equipment and data-flow schematic for the multi-modal experiment.

### 2.3. Task and Stimulus

Tasks involved the visual sorting of five-digit numbers as they appeared on a computer screen. The numbers resembled the postal codes used in the United States (Figure 2). Participants were asked to match a randomly generated postal code (shown in red in the middle of the screen) to its corresponding range (shown on the left of the screen). There were a total of six ranges, each identified by a color that corresponded to the colors of the buttons on the Cedrus response box. The correct range was indicated by pressing the corresponding color-coded button. A pie-chart marking the progress of the experiment was displayed on the right of the screen. The number of correct responses and number of remaining trials were provided as feedback to the participants. Participants were guided to perform their tasks by sequential instructions shown on the screen, and to provide their responses as required during stimulus presentation. EEG, GSR, and eye-movements were monitored continuously. Stimulus and responses along with other events were time-stamped.

**Figure 2:**
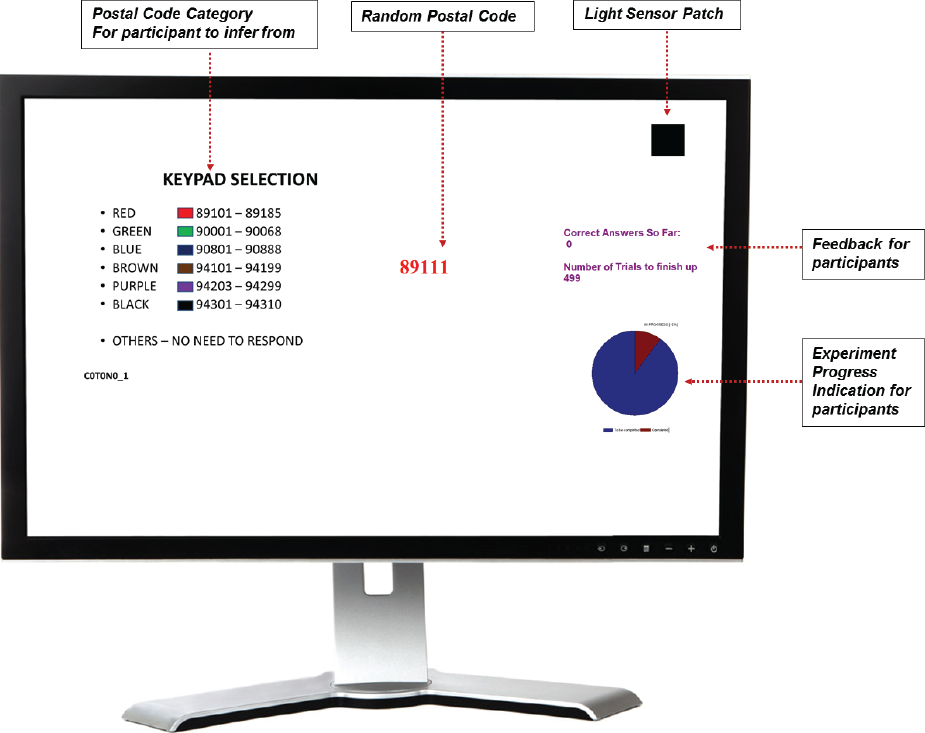
Screenshot with stimuli presented to the participant in one trial.

### 2.4. Experiment Design

A sequential series of tasks was designed to induce increasing levels of task difficulty by manipulating three binary-valued variables: Color (C), Numerical arrangement of the six ranges (N), and Time (T). The CNT triplet of binary values yields 8 possible values, each constituting a block. In each block, numeric “postal” codes were presented 40 times without repetition. Within a block, *C* = 0 if the color of each range is held constant across trials; *C* = 1 if the color for each range is randomly shuffled for every trial. *N* = 0 if the arrangement of the range labels on the screen does not change across trials, whereas *N* = 1 if the range labels are scrambled every trial. *T* = 0 if the allotted response-time is kept constant (at 7 seconds) and *T* = 1 if the allotted response-time is variable (chosen randomly from the interval 2-7 seconds with uniform probability). We hypothesize that when task conditions are changed (i.e., when C, T, or N are 1) the task load increases. Thus, the easiest task is CTN = 000 (the first block in the sequence of tasks, labeled CTN000), and the most difficult is CTN = 111 (the last block in the sequence of tasks, labeled CTN111). The last block was repeated (CTN111A), resulting in 9 blocks and 360 trials in total. The time sequence of the trials is depicted in Figure 3.

**Figure 3:**
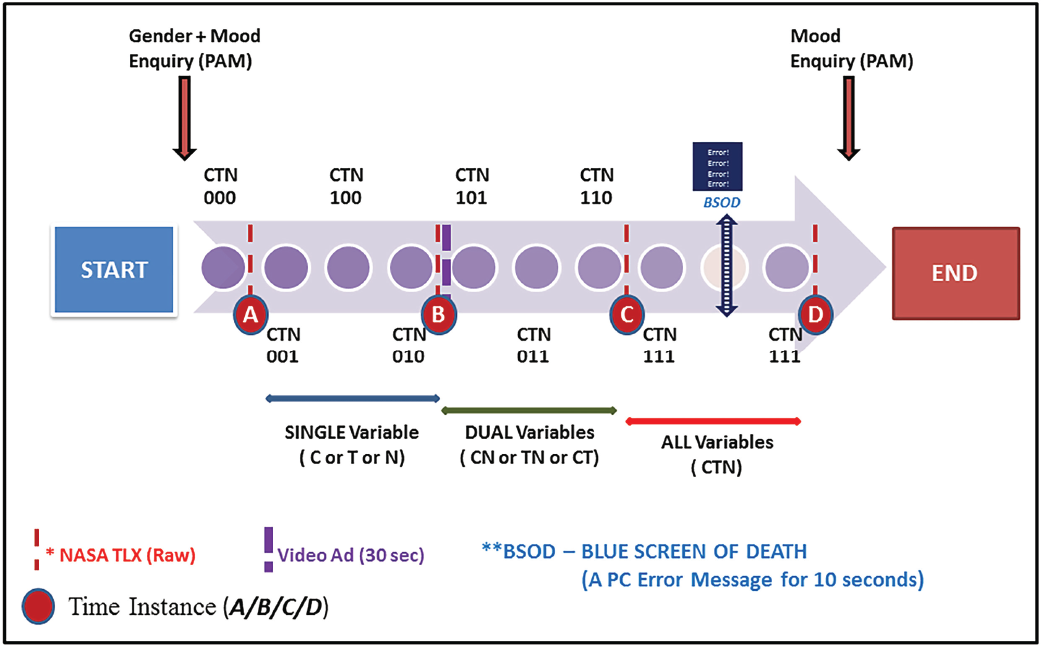
Protocol depicting the time sequence of the experiment.

The task flow is interrupted at several time points in between blocks. At time points A-D (see Figure 3), the participant is asked electronically to respond to the NASA-TLX questionnaire (responses are time-stamped). At time point B, the tasks are interrupted and the participant is required to watch a video (of a randomly selected advertisement) for 30 seconds. Between time points C and D, the blocks CTN111 and CTN111A are separated in time by the appearance of a “Blue screen of death” (or BSOD, a PC error message) for 10 seconds. These two events were hypothesized to cause a subjective and objective increase in task load in the block immediately following the event.

### 2.5. Grouping of Data

The blocks CTN000 to CTN111, with a repeat CTN111A, can be tested for across-block differences in task difficulty. However, it is possible that some combinations of the blocks may have similar levels of difficulty. Therefore, to improve the discrimination of task difficulty, we grouped together blocks based on properties that are assumed to be shared. While many such groupings are possible, we decided on eight groups shown in Table 1. These are:

- Rows 1-4: Groupings considering single variables C, T, N, or the temporal flow of the experiment. Row 4 (*Temporal Flow*) is the most fine-grained grouping retaining all 8 possible values.
- Rows 5-8: Groupings considering the hypothetical difficulty of the task. Row 5 groups the interaction of the C and N variable (which differentiates blocks in which visual search was required to solve the task, i.e. *C* = 1 ∨ *N* = 1, from blocks with steady legends). Row 6, *Hypothetical Difficulty*, groups the modulation of task difficulty by the binary task variables. Row 7, *# Variables* (number of variables) groups blocks which have the same number of manipulations of C, T, and N. Row 8, < 2 *Variables*, groups manipulated variables together to reflect the participant’s adaptation to the task. Here, groups that are 0 and 1 in row 7 are grouped as 0, group 2 in row 7 is now group 1, and group 3 is 2.

**Table 1:** Grouping of blocks, based on combinations of the C, T, and N variables, to reduce the number of manipulated variables. Each row depicts a group of blocks. Groupings depend on the manipulation of the 3 binary variables, temporal flow, or hypothetical task difficulty.

| Row | Grouping | 000 | 001 | 100 | 010 | 101 | 011 | 110 | 111 |
| --- | --- | --- | --- | --- | --- | --- | --- | --- | --- |
| 1 | Color (C) | 0 | 0 | 1 | 0 | 1 | 0 | 1 | 1 |
| 2 | Time (T) | 0 | 0 | 0 | 1 | 0 | 1 | 1 | 1 |
| 3 | Postal Code (N) | 0 | 1 | 0 | 0 | 1 | 1 | 0 | 1 |
| 4 | Temporal Flow | 1 | 2 | 3 | 4 | 5 | 6 | 7 | 8 |
| 5 | C+N | 0 | 1 | 1 | 0 | 1 | 1 | 1 | 1 |
| 6 | Hypo. Diff. | 0 | 1 | 1 | 2 | 2 | 3 | 3 | 4 |
| 7 | # Variables | 0 | 1 | 1 | 1 | 2 | 2 | 2 | 3 |
| 8 | <2 Variables | 0 | 0 | 0 | 0 | 1 | 1 | 1 | 2 |

Finally, a *post-hoc* grouping of the *individual performance* was done (not shown in Table 1). Based on the standardized intra-subject performance *z*, blocks were classified into one of the three classes *low* (*z* < −0.6), *medium* (−0.6 < *z* < 0.6), *high performance* (*z* > 0.6).

In the single-modality analysis we used the data from individual modalities (GSR, EEG, Eye-tracking) and searched for significant differences between the groups using one-way ANOVA. We tested for *H*_0_: *μ*_1_ = *μ*_2_ *·*= *μ*_*i*_ against *H*_1_ where there are at least 2 groups of means. In case ANOVA results revealed significance, post-hoc Tukeys Honest Significant Difference (HSD) tests (multiple comparison test between all combinations of groups) were conducted.

## 3. Data Processing

### Software

All data were analyzed using Matlab (The MathWorks, Inc). EEG data were analyzed using EEGLAB (Delorme and Makeig, 2004), GSR data with Ledalab (Benedek and Kaernbach, 2010a,b); both are open-source toolboxes for Matlab. Eye-tracking data were processed by SMI BeGaze, integrated with the SMI Experiment Center and analyzed in Matlab.

### Eye Tracking

Data were averaged within each block of trials, and then standardized across all blocks within each participant.

### Galvanic Skin Response (GSR)

GSR data were analyzed in terms of phasic and tonic skin conductance components (in *μS*) after pre-processing (down-sampling to 10 Hz, filtering with a 4th-order IIR filter having cutoff frequency 2 Hz, smoothing with moving average window of 100 samples, and segmenting data using the event triggers generated by the recording system). GSR feature values were calculated for every subject and condition as a mean response of all trials, and converted to standardized scores.

### Eye Tracking

Data were averaged within each block of trials, and then standardized across all blocks within each participant.

### Electroencephalography (EEG)

EEG data were acquired at 2500 Hz from 32 electrodes. The following preprocessing pipeline was applied to the raw data, based on the detrending procedure suggested by Cohen (2014).

1. **Removal of Line Noise**: A notch filter was applied in order to remove the 50 Hz frequency component and its harmonics up to 250 Hz.
2. **Epoching**: To facilitate the study of task-related changes, the continuous data were cut into time segments surrounding the events. The epochs were defined within a time window of [−1.5; +1.5] seconds from the onset at time 0 (the appearance of the postal code on the screen).
3. **Channel rejection** of electrodes that showed artifacts or a mean channel power over three standard deviations from the mean among all channels. This resulted in the rejection of the electrodes Fp, FPz and Fp2 due to eye blink contamination, and M1 and T7 due to the presence of EMG artifacts, leaving 27 electrodes.
4. **Spatial filter**: Application of a common average removal (CAR) spatial filter was chosen as an alternative to other spatial filters that improve resolution to localized sources, due to the spread nature of the measured EEG response.
5. **Filtering**: Application of a 4^th^-order finite impulse response (FIR) bandpass filter, preserving a frequency band from 0.5 to 80 Hz.
6. **Rejection of epochs** containing an abnormally larger power in comparison to other epochs, a linear drift in the signal or movement artifacts.
7. **ICA for artifact rejection**: Applying the logistic infomax ICA decomposition (Bell and Sejnowski, 1995) on the epoched data as an artifact removal method in order to reject components that were likely to be caused by blink artifacts (strong frontal activation, steep power spectrum) and/or muscular activity (spatially localized activity, high power above 20 Hz).
8. **Normalization**: A baseline referencing of each data epoch on a time window of [−1.5;−0.1] seconds from stimulus onset was chosen with a window of the same length as the epochs (±1.5 seconds) to compute the Welch power spectral density (PSD) (Welch, 1967). This choice is based on the PCA decomposition of the PSD data, which showed that this normalization approach resulted in the largest proportion of variance (71%) being explained by the first three components.
9. **Feature extraction**: The EEG features were extracted in the frequency domain via PSD with Welch’s overlapped segment averaging estimator. This computes a modified periodogram for each segment window and then averages these estimates to produce the estimate of the PSD. As opposed to the standard periodogram, it reduces noise with the trade-off of having lower frequency resolution. Its Hamming window further prevents ripple effects at window extremities. For this method, the epochs were divided into sliding windows of 500ms with an overlap of 50% between each other, as suggested by Zhang et al. (2014). The PSD was calculated inside a range of 0.5 to 45 Hz, taking 30 equally distributed frequency bins.

The initial feature extraction step results in 810 features (30 frequency bins for 27 electrodes). Such a large number of features creates a computationally expensive classification problem. Furthermore, it results in estimation errors due to reduced sample size (for each feature). Thus, we resort to dimension reduction so as to increase the sample size and estimation accuracy, as follows.

1. **Grouping of frequency bins into frequency bands:** The mean of the PSD was calculated for the frequency bins in the following 8 bands: Delta (0.5-3.5 Hz), Theta (3.5-7.5 Hz), Alpha (7.5-12.5 Hz), low-Beta (12.5-18 Hz), mid-Beta (18-24 Hz), high-Beta (24-30 Hz), low-Gamma (30-37.5 Hz), and Gamma (37.5-45 Hz), resulting in 216 features (one value per band and electrode).
2. **Preselection of electrodes:**Features were sorted by the Fisher Score, which ranks the features so as to maximize the distances between data points from different groups and minimizes the distances between data points in the same class. It is defined as:

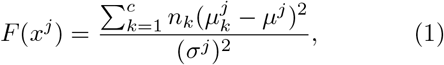

where 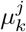 and 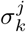 are the mean and standard deviation of the *k*^th^ group or class respectively, corresponding to the *j*^th^ feature; *μ*^*j*^ and *σ*^*j*^ the mean and standard deviation of the whole data set corresponding to the *j*^th^ feature, *n*_*k*_ the size of the *k*^th^ class and 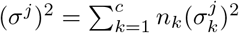.
3. **ANOVA tests of the remaining electrodes:**The trials were grouped into classes depending on the *# Variables* grouping (Table 1, row 7). Next, we selected the electrodes that were represented in the 100 features with the highest Fisher score. The final set of electrodes and frequency bands were determined from the results of the ANOVA test for each pre-selected electrode in each frequency band. For each location group (see Table 2), the electrode with the most significant ANOVA tests for the previously described grouping was chosen, so as to have a uniform distribution of the tested areas. The selection of the frequency bands for analysis was based on the same criterion.

**Table 2:** Electrodes in spatial proximity were grouped into 7 location groups. From every group, one electrode (in bold) was picked for posterior analysis.

| Location Group | Electrode Number | Electrode Names |
| --- | --- | --- |
| A | 5, 9, 10, 15 | F3, FC5, **FC1**, C3 |
| B | 7, 11, 12, 17 | F4, FC2, **FC6**, C4 |
| C | 15, 20, 21, 25 | C3, CP5, **CP1**, P3 |
| D | 17, 22, 23, 27 | **C4**, CP2, CP6, P4 |
| E | 10, 11, 16, 21, 22 | FC1, FC2, **Cz**, CP1, CP2 |
| F | 5, 6, 7 | F3, Fz, **F4** |
| G | 29, 30, 31, 32 | **POz**, O1, Oz, O2 |

This approach resulted in the selection of the following 7 electrodes: FC1, FC6, CP1, C4, Cz, F4, POz. The 6 most significant frequency bands were Alpha, low-Beta, mid-Beta, high-Beta, low-Gamma, and Gamma, resulting in 42 EEG features for the posterior analysis.

## 4. Results

### 4.1. Behavioural Responses

Figure 4 shows the fraction of correct responses as a function of the nine blocks (the permutations of the values taken by the variables C, T, and N, hereafter referred to as CTN blocks to distinguish them from other blocks depicted in Table 1). The blocks are arranged along the abscissa in their temporal order of presentation. The proportion of correctly sorted postal codes degrades over time (i.e., blocks), with a clear separation of the first 5 blocks from the last 4. As hypothetical task difficulty increases, the rate of correct answers declines.

**Figure 4:**
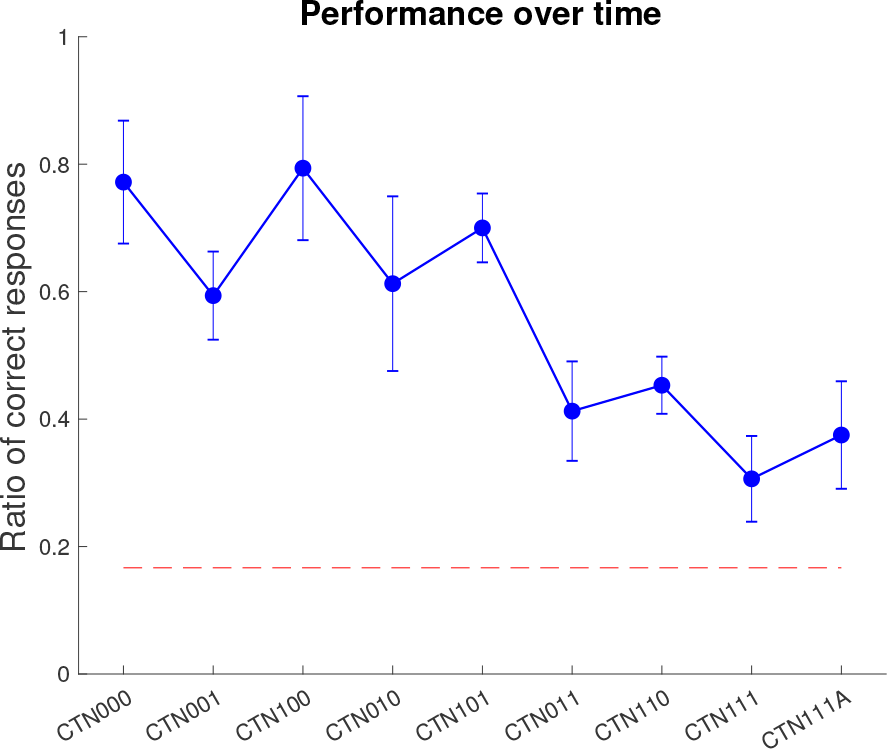
Fraction of correct responses averaged across participants for each block, where C, T, and N assume binary values (CTN111A is a repeat of CTN111). The responses are arranged in the sequential order of presentation. Bars represent 95% confidence intervals, the dashed red line indicates performance at chance level (16.67%).

For each CTN block, Figure 5 depicts the average response time of correct and incorrect responses. Across all blocks, incorrect responses were quicker (smaller response times) than correct responses.

**Figure 5:**
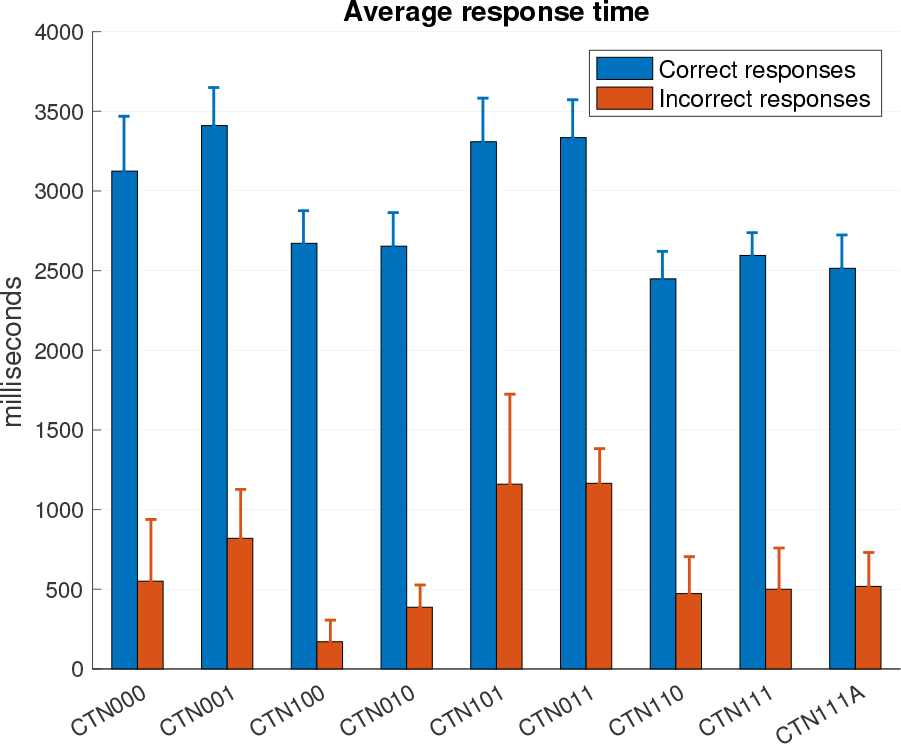
Average response time for both correct and incorrect responses. Error bars represent 95% confidence intervals. Quicker responses were more likely to be incorrect.

Figure 6 shows the z-scores for error (inverse of the performance) and response time as a function of increasing number of manipulated variables C, T, and N (Table 1, row 7, *# Variables*). Blocks are grouped together if they have the same number of manipulated variables. The assumption is that the level of task difficulty increases as more variables are manipulated (i.e., going from left to right along the abscissa). Multiple comparison of all difficulty pairs (0 vs. 1, 1 vs. 2, 2 vs. 3) yields statistically significant differences. Response time on the other hand does not vary significantly across groups, and is almost uncorrelated (*r* = *−*0.07) with task performance.

**Figure 6:**
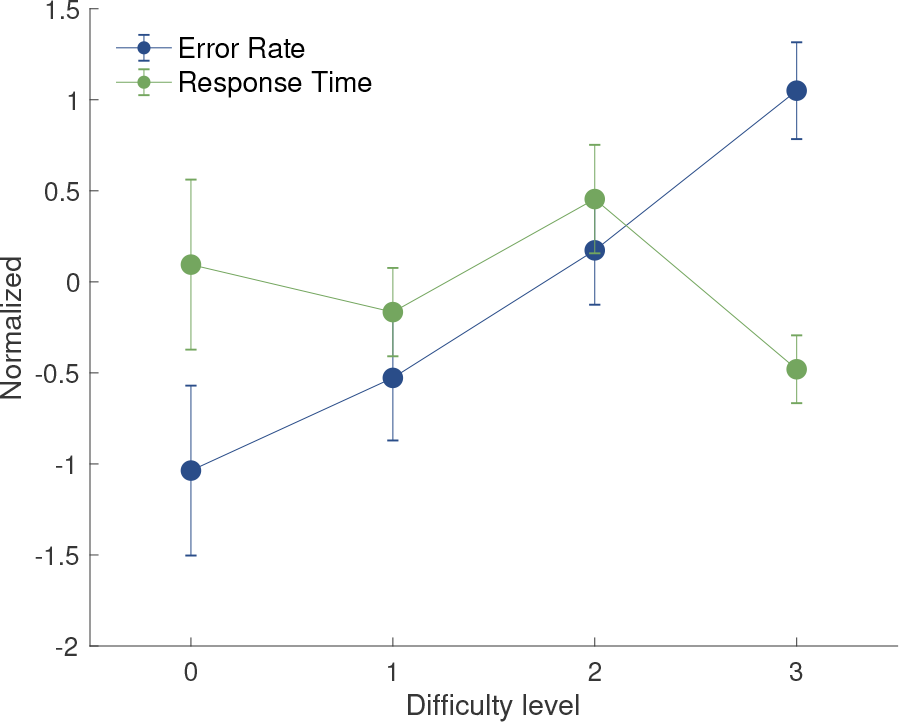
Interval plot of z-scores for error (blue, inverse of the performance), and response time (green), as a function of hypothetical task difficulty.

### 4.2. NASA TLX Questionnaire

At various points in time throughout the experiment (denoted A, B, C and D in Figure 3) participants were asked to provide a self-assessment of the perceived level of workload based on the NASA TLX instrument. This psychometric assessment comprises six aspects, namely performance, effort, frustration, as well as mental, physical, and temporal workload. For each of these aspects, ratings were collected on a 7-point Likert scale, from 1 (very low) to 7 (very high). The grand averages of individual ratings are depicted in Figure 7. All six aspects had the lowest scores at the first time point (easiest task) and increased thereafter, albeit at different rates.

**Figure 7:**
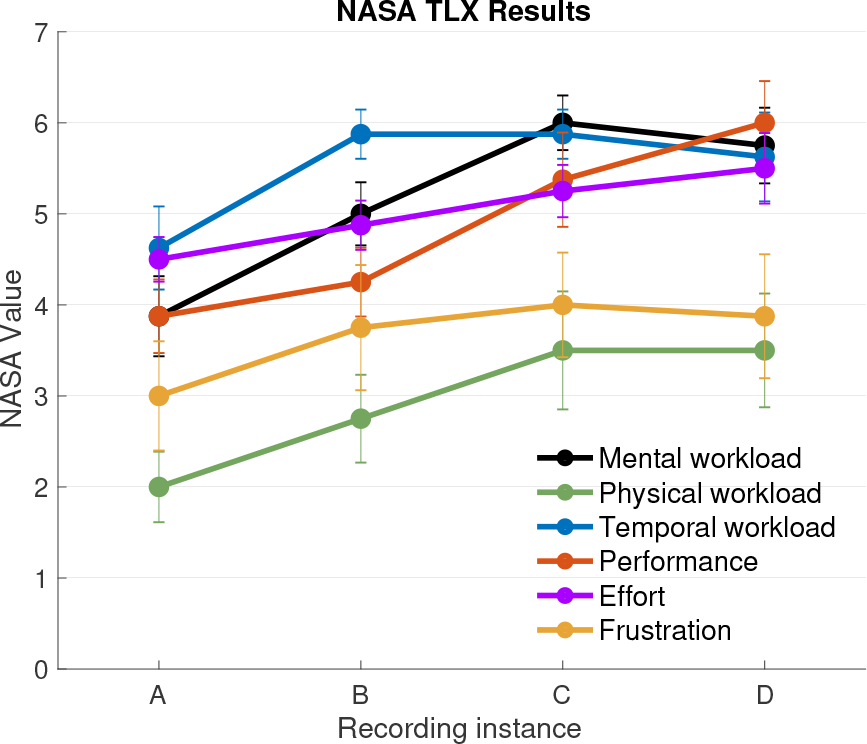
NASA Task Load Index ratings (self-assessment) were collected 4 times during the experiment (A-D). Each data point is the grand average of the ratings across participants. Error bars depict confidence intervals.

### 4.3. EEG

Figure 8 shows representative EEG images of the most active regions on the scalp for a set of 6 groupings. Activity was most prevalent in central and parietal regions of the right hemisphere and strongest for the initially assumed grouping of task difficulty (Table 1, row 6, *Hypothetical Difficulty*; Figure 8, bottom row, center) – the only grouping in which a tendency for symmetric activity could be revealed. Activity in frontocentral regions (particularly electrode FC6) underlies modulation by task performance.

**Figure 8:**
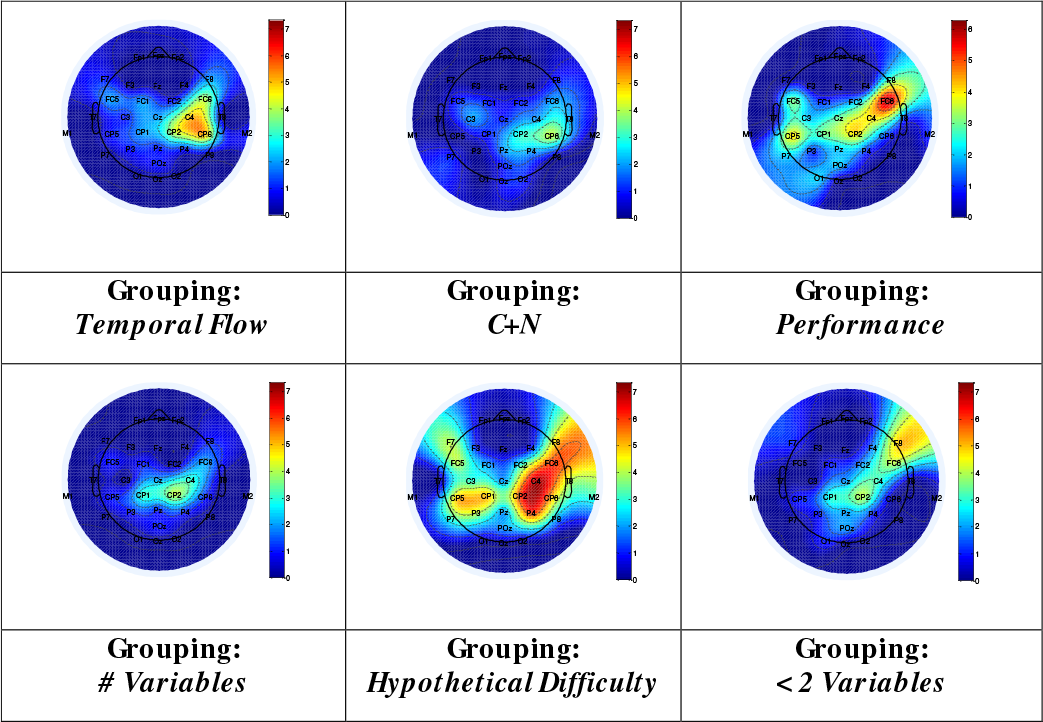
EEG topoplots depicting for how many participants a certain channel showed a significant response (*p* < 0.05) based on the indicated groupings.

In the following, we outline the results of the ANOVA tests for the selected electrodes, frequency bands and groupings.

#### Grouping by *Temporal Flow*

This group is depicted in row 4, Table 1, and reflects the chronological flow of time along with a gradual increase in task difficulty. One-way ANOVA test results for significant response are shown in Table 3. Although the power in the low-Beta band for FC6 was itself not significant, the low-Beta band had the most significant response when viewed across the selected electrodes, whereas FC6 had the most significant response across frequency bands.

**Table 3:** One-way ANOVA test of EEG signal power in various frequency bands and electrodes when grouped according to *Temporal Flow* (CTN blocks 1 to 8, see row 4, Table 1). Significant *p*-values are highlighted in red (*p* < 0.001) and pink (0.001 < *p* < 0.05).

| Electrode | Alpha | Low Beta | Mid Beta | High Beta | Low Gamma | Gamma |
| --- | --- | --- | --- | --- | --- | --- |
| 10/FC1 | 0.83 | 3E-3 | 0.04 | 0.20 | 0.53 | 0.13 |
| 12/FC6 | 0.07 | 0.09 | 0.03 | 0.01 | 0.01 | 0.01 |
| 21/CP1 | 0.20 | 2E-4 | 0.09 | 0.11 | 0.07 | 0.03 |
| 17/C4 | 0.01 | 0.04 | 0.09 | 0.05 | 0.04 | 0.02 |
| 16/Cz | 0.77 | 0.01 | 0.04 | 0.32 | 0.49 | 0.21 |
| 7/F4 | 0.27 | 0.05 | 0.09 | 0.30 | 0.26 | 0.33 |
| 29/POz | 0.23 | 6E-4 | 0.17 | 0.27 | 0.57 | 0.21 |

#### Grouping by *Time (T)*

This group is depicted in row 2, Table 1 and reflects the manipulation of time. One-way ANOVA test results for significant response are shown in Table 4. Responses were most significant in the three Beta bands (especially low-Beta) across most electrodes, and for the FC6 electrode across all frequency bands. In all cases the PSD features increased in their standardized scores with the change from *T* = 0 to *T* = 1.

**Table 4:** Grouping by *Time (T)* (see row 2, Table 1). Description follows Table 3.

| Electrode | Alpha | Low Beta | Mid Beta | High Beta | Low Gamma | Gamma |
| --- | --- | --- | --- | --- | --- | --- |
| 10/FC1 | 0.82 | 9E-4 | 0.09 | 0.25 | 0.52 | 0.31 |
| 12/FC6 | 5E-3 | 8E-4 | 4E-3 | 0.02 | 0.02 | 6E-3 |
| 21/CP1 | 0.08 | 8E-4 | 4E-3 | 0.01 | 0.19 | 0.15 |
| 17/C4 | 0.01 | 2E-3 | 0.02 | 0.04 | 0.12 | 0.08 |
| 16/Cz | 0.16 | 4E-4 | 2E-3 | 0.02 | 0.16 | 0.20 |
| 7/F4 | 0.31 | 2E-3 | 0.01 | 0.04 | 0.06 | 0.08 |
| 29/POz | 0.11 | 8E-4 | 0.01 | 0.07 | 0.23 | 0.17 |

#### Grouping by *Hypothetical Difficulty*

This group is depicted in row 6, Table 1, and reflects increasing task difficulty as variables (C, T, N) are manipulated, with additional weight given to the manipulation of the time (T) variable. One-way ANOVA test results for significant response are shown in Table 5. One-way ANOVA tests of the remaining groupings (rows 1, 3, 5, 7, and 8 in Table 1) did not reveal any significant responses over the range of electrodes or frequency bands and are therefore not presented here. To obtain Figure 9, the error rates of all participants in all blocks were standardized per subject and then correlated with the PSD of a particular channel at a specific frequency band. Consistently across the presented electrodes and frequency bands (excluding alpha), increased error rate was accompanied by increased PSD values.

**Table 5:**
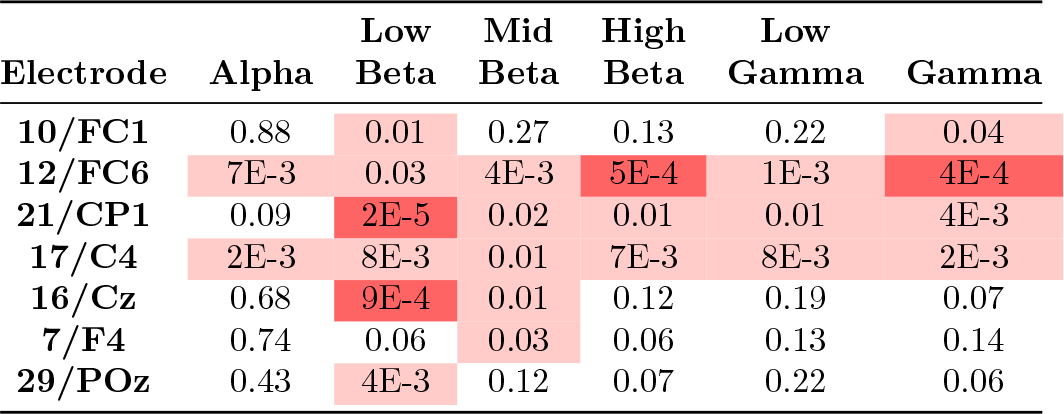
Grouping by *Hypothetical Difficulty* (see row 6, Table 1). Description follows Table 3.

| Electrode | Alpha | Low Beta | Mid Beta | High Beta | Low Gamma | Gamma |
| --- | --- | --- | --- | --- | --- | --- |
| 10/FC1 | 0.88 | 0.01 | 0.27 | 0.13 | 0.22 | 0.04 |
| 12/FC6 | 7E-3 | 0.03 | 4E-3 | 5E-4 | 1E-3 | 4E-4 |
| 21/CP1 | 0.09 | 2E-5 | 0.02 | 0.01 | 0.01 | 4E-3 |
| 17/C4 | 2E-3 | 8E-3 | 0.01 | 7E-3 | 8E-3 | 2E-3 |
| 16/Cz | 0.68 | 9E-4 | 0.01 | 0.12 | 0.19 | 0.07 |
| 7/F4 | 0.74 | 0.06 | 0.03 | 0.06 | 0.13 | 0.14 |
| 29/POz | 0.43 | 4E-3 | 0.12 | 0.07 | 0.22 | 0.06 |

**Figure 9:**
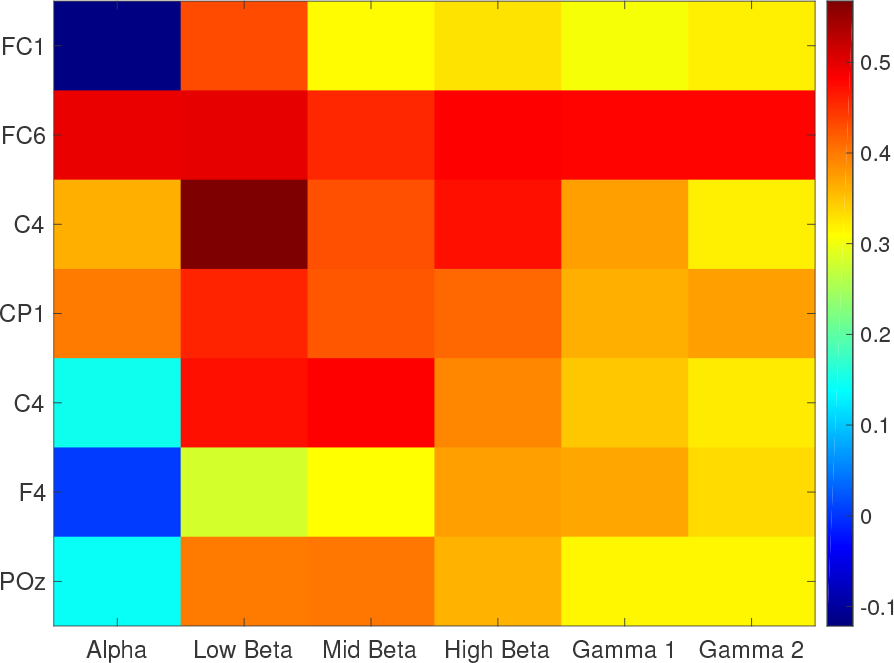
Pearson correlation coefficient of the seven preselected electrodes with the individual error rates. Correlations are shown for each frequency band individually.

#### EEG Asymmetry Index

The EEG asymmetry index quantifies hemispherical imbalances of cortical activity and was calculated to investigate the apparent greater responsiveness of the FC6 electrode compared to its contralateral equivalent (FC5):

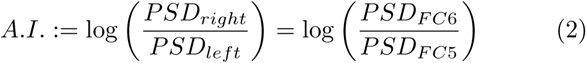

ANOVA tests of the asymmetry index on the change between conditions of low and high workload revealed significances for all tested groupings, in particular for the *# Variables* grouping (*p* < 0.005). Figure 10 depicts the increase in asymmetry index for levels of higher difficulty.

**Figure 10:**
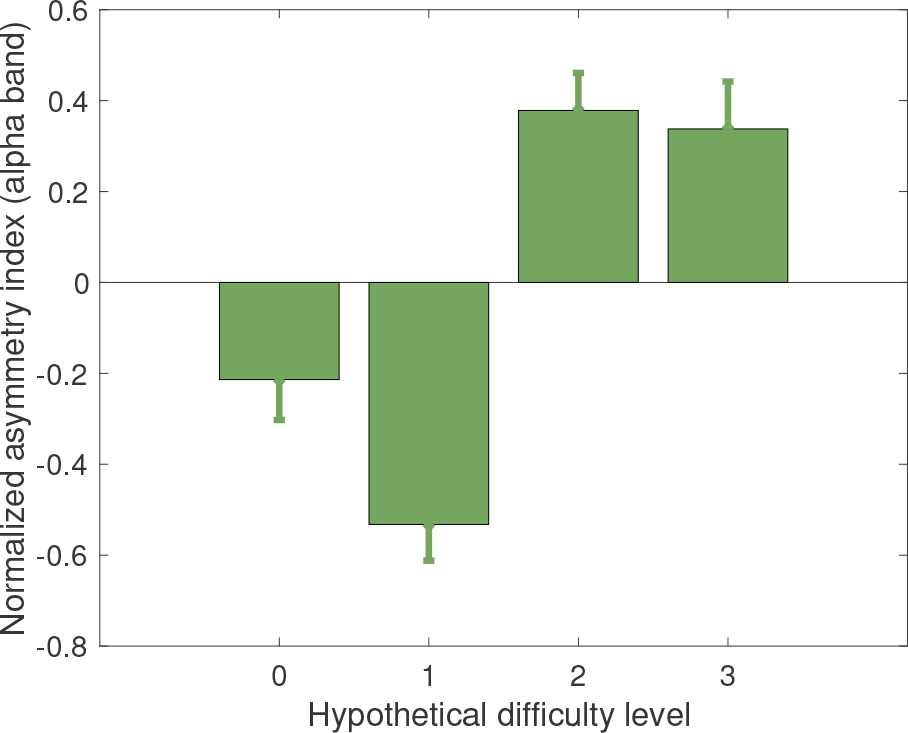
The EEG-asymmetry index of the electrodes FC6 and FC5 reveals an increase in fronto-central regions of the right hemisphere as a function of cognitive load.

### 4.4. Galvanic Skin Response

The Galvanic Skin Response (GSR) is usually decomposed into two components: i) the tonic response (slow changes in the GSR signal with time-scales ranging from seconds to minutes), also called Skin Conductance Level (SCL), and ii) the phasic response (rapid changes in the GSR signal with time-scale up to seconds), named Skin Conductance Response (SCR). Results of one-way ANOVA tests are shown in Table 6, for four groups taken from Table 1 (rows 2, 5, 6, and 7). Due to the high correlation between various SCR- and SCL-related features (the full list of tested features is available in the Appendix), only the results for mean amplitude of both components are presented.

**Table 6:** *p*-values of one-way ANOVA for representative GSR features across selected groupings. Pink indicates 0.001 < *p* < 0.05, and red indicates *p* < 0.001.

| Grouping | SCR amplitude | SCL amplitude |
| --- | --- | --- |
| Time (T) | 0.02 | 0.59 |
| C+N | 0.05 | 0.65 |
| Hypo. Diff. | 9E-5 | 0.06 |
| # Variables | 7E-5 | 0.02 |

### 4.5. Eye Tracking Data

Of the eleven eye-tracking features tested (see Appendix for a complete list), we focus on fixation duration, fixation positions, blink duration, and pupil diameter, which were found to be correlated with task difficulty and cognitive load, and can potentially determine blocks where performance exceeds chance level. There were no significant findings for distance to the screen, ratio and duration of saccades.

#### Fixation Duration

Average duration of fixations in our data showed an 11% reduction in fixation (*p* < 3e−7) in the blocks that required visual search for solving the task (*C+N* grouping). Although the participants had to solve the same sorting task in principle, they fixated much longer on a position on the screen if visual search was not required. Thus, a task requiring visual search involves shorter fixation durations than the same task without visual search.

#### Fixation Positions

Fixations of gaze on the screen were analysed in detail by generating heatmaps so as to visualize the areas of visual interest (see Figure 11). In general, participants rarely read the names of the colors (shown on the extreme left of the screen), but perceived the color button peripherally while fixating on the lower bound of the postal code range (the upper bound was mostly ignored). Next, fixations on the rightmost digits of the 5-digit postal code diminished over time, since participants realized their irrelevance for the decision (both for the current postal code (red) shown in the center of the screen, and the legend shown to the left of the screen). Furthermore, fixation positions were more scattered in later blocks, potentially indicating a less goal-oriented behavior in more difficult tasks.

**Figure 11:**
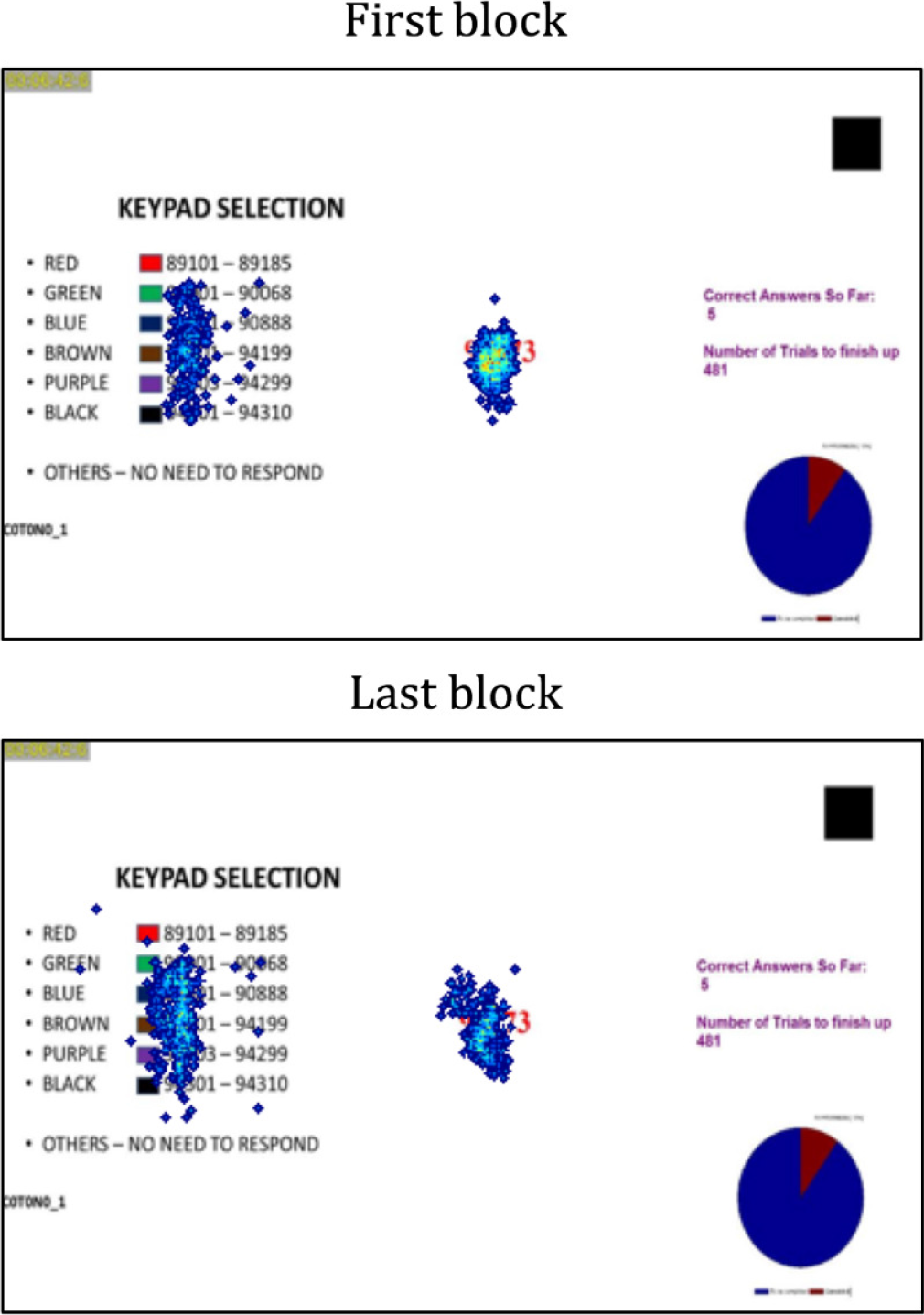
Position of most frequent fixations (thresholded) for the first block (top) and last block (bottom). Heatmaps show averages across all participants.

In order to quantify this visually inferred tendency, we trained a simple multi-layer perceptron (MLP) to predict the position of the next fixation point based solely on the position of the current fixation point. Assuming that a frustrated subject would be more likely to get distracted from the task and randomly perform saccades while viewing the screen, we hypothesized that for blocks with worse performance the classification accuracy of the MLP would diminish. The classification was based on a discretization of the screen into bins of equal size, and the fixation positions of all agents were used as training data. As hypothesized, higher predictability of the fixation positions in one block (the accuracy of the MLP) was accompanied by better performance. A blockwise comparison of (standardized) network accuracy and performance revealed a high correlation, supporting our assumption that more random or unexpected fixations occurred in more challenging blocks (*r* = 0.59, see Figure 12). In other words, a more structured, goal-oriented behavior in blocks with better performance could be inferred from the data. A hypothesis test comparing the predictability of fixation positions in blocks above and below average performance was significant (*p* < 4e*−*5).

**Figure 12:**
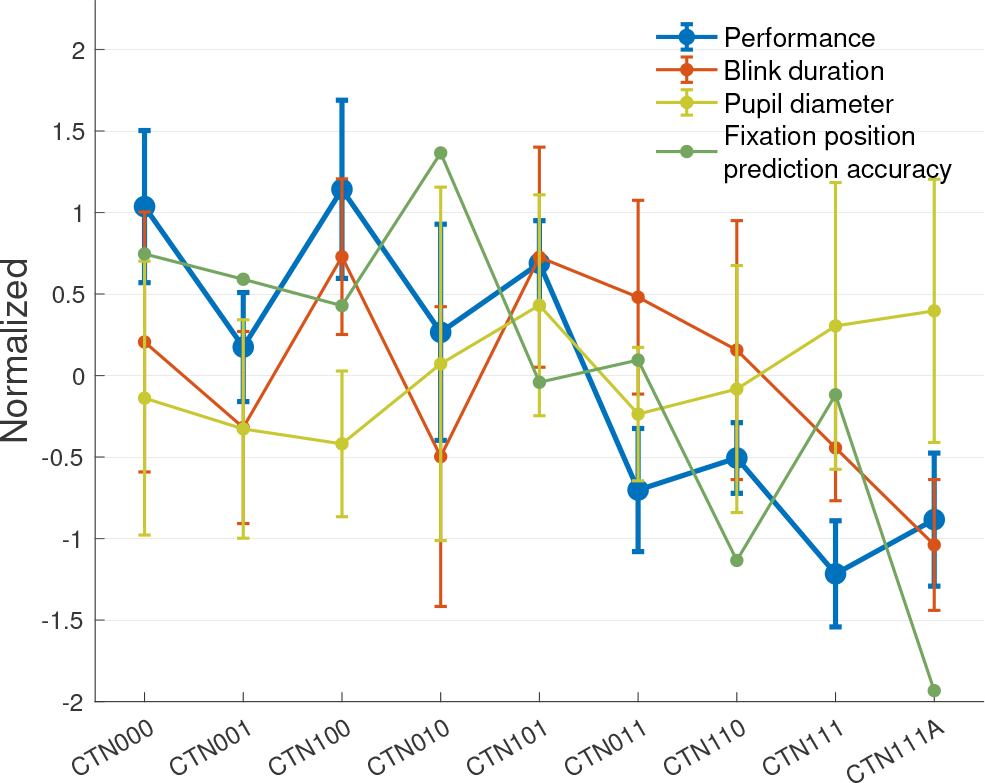
A blockwise comparison of task performance with blink duration and fixation position predictability revealed high correlations (*r* = 0.57 and *r* = 0.59) and a weaker, negative correlation with pupil diameter (*r* = −0.4). Thus, participants had less structured fixation trajectories in blocks of lower performance.

#### Blink Duration

Blink duration is thought to be a reliable measurement of drowsiness. Our data shows positive correlation between blink duration and task performance (*r* = 0.57, see Figure 12). For our data, the ANOVA test allows distinctions based on the individual performance, i.e., between blocks with satisfactory performance (standardized score > 0.5) and those with poor performance (standardized score < −0.5, post-hoc Tukeys HSD test: *p* = 0.001), and medium performance (*p* = 0.03).

#### Pupillometry

Since pupil diameter is mostly sensitive to variation in illumination rather than cognitive states (Dehais et al., 2008), we conservatively removed the 10 seconds of data following the inter-block pauses that were interrupted by distractions (see Figure 3). The remaining measurements indicate an increase in pupil diameter with task difficulty (*r* = 0.63*, p* < 0.02), confirming previous findings.

### 4.6. Multimodal Analysis

Multimodal analysis attempts to quantify the amount of information carried by the features of the previously examined modalities and determines which physiological measures are reliable. For example, do we gain additional information from eyetracking data in combination with EEG?

Perhaps the most direct method is to employ a multimodal model that iteratively adds independent variables from the pool of features based on some optimization criterion. From the various regression models we chose a linear model (implemented in Matlab via fitlm and step) due to its simplicity and interpretability (it can quantify the relevance for performance prediction for each of the physiological measurements and their individual features).

First, linear regression models were computed for all three modalities (EEG, GSR, and ET, i.e., eye-tracking) separately (top 3 rows in Table 7). Models started with all features and improved stepwise by adding or removing terms based on the *p*-value of an F-test as an optimization criterion.

**Table 7:** Linear Regression models were computed from the features of individual modalities (top 3 rows) and various combinations. Model quality was assessed by RMSE and adjusted R^2^. The full model outperforms all others. For a legend of the feature indices, see the Appendix.

| Modalities | Details | RMSE | $R^2$ | Adj. $R^2$ | p-value | # | Features |
| --- | --- | --- | --- | --- | --- | --- | --- |
| **GSR** | Stepwise | 0.81 | 0.32 | 0.28 | 1E-04 | 4 | GSR: 1,2,5,15 |
| **ET** | Stepwise | 0.79 | 0.36 | 0.30 | 9E-05 | 5 | ET: 3,4,8,10,11 |
| **EEG** | Stepwise | 0.56 | 0.74 | 0.65 | 8E-09 | 16 | EEG: 1-4,6,10,14-17,20,26-28,31,34 |
| **GSR+ET** | Pre-selected | 0.72 | 0.51 | 0.43 | 8E-06 | 9 |  |
| **GSR+ET** | Step+Pre | 0.72 | 0.49 | 0.42 | 3E-06 | 7 | GSR: 1,2 ET: 3,4,8,10,11 |
| **EEG+ET** | Pre-selected | 0.55 | 0.78 | 0.66 | 1E-07 | 21 |  |
| **EEG+ET** | Step+Pre | 0.55 | 0.73 | 0.66 | 6E-10 | 13 | EEG: 1-4,6,10,14,17,26,27,34 ET: 4,11 |
| **EEG+GSR** | Pre-selected | 0.50 | 0.81 | 0.72 | 2E-09 | 20 |  |
| **EEG+GSR** | Step+Pre | 0.51 | 0.79 | 0.71 | 2E-10 | 13 | EEG: 1-4,6,10,14-17,26,27,31,34 GSR: 1,2 |
| **ALL** | Pre-selected | 0.52 | 0.80 | 0.72 | 1E-07 | 25 |  |
| **ALL** | Step+Pre | 0.50 | 0.80 | 0.72 | 2E-10 | 16 | EEG: 1-4,6,10,14-17,26,27,31,34 ET: 4 GSR: 1,2 |

For combinations of modalities, the features selected on the single-modality models were used (pre-selected) before the same optimization technique was executed (step+pre). For model comparison, root mean squared error (RMSE), the adjusted *R*^2^ (i.e. the squared correlation coefficient weighted by the number of features used in the model) and the mentioned *p*-value of the model vs. a constant model (guessing the mean) were considered. Since RMSE is a scale-dependent parameter, all variables were normalized. The constant model yielded an RMSE of 0.935.

The results in Table 7 suggest that EEG data is by far the most informative to infer about task performance, compared with eyetracking and GSR data. Even combining the latter two modalities yields worse results than EEG alone. Within the single-modality models, the most discriminant features of the ANOVA analysis (e.g. blink duration in eyetracking) had the highest coefficients (in absolute terms) in the linear model. As expected, the EEG models were dominated by features of the low Beta band, in particular fronto-central electrodes (compare Figure 9). Importantly, the full model (taking into consideration all modalities) outperforms all others, although the *EEG+GSR* model comes very close. A repetition of model fitting with a 10-fold cross validation, conducted in an attempt to obtain a more robust estimate of model quality, yields nearly the same values as in Table 7.

## 5. Discussion

### 5.1. Behavioural Responses

We noticed a clear degradation in percent-correct responses with increase in time (Figure 4), while there was also a concomitant increase in hypothesized task difficulty. This may suggest that (1) task difficulty indeed increases with time, and (2) that a joint manipulation of the time variable and at least one of the legend variables (C, N) severely hampers sorting performance. Presumably, performance was subject to online learning processes, as indicated by the increases between block pairs (CTN001, CTN100), (CTN011, CTN110) and (CTN111, CTN111A) where the alteration in task difficulty was minimal or absent.

The perceived level of workload was assessed from the NASA TLX questionnaire at four time points (Figure 3) for six aspects, namely performance, effort, and frustration, and for mental, physical, and temporal workload (Figure 7). All six aspects had the lowest scores at the first time point (easiest task) and thereafter increased, albeit at different rates. This suggests that the loading of the task had a differential effect on each aspect. Unfortunately, the sample size was too small for a more in-depth statistical analysis, so we provide only a qualitative description of the results.

Physical workload had the lowest scores overall across all time-points, as the task made almost no physical demands. However, physical workload scores showed an increase when two and three (all) variables were changed but demonstrated larger standard deviations, reflecting perhaps an inability to appropriately scale the perceived physical effort (i.e., the noise in the estimate dominates any significant change in the score). This was also true for frustration which demonstrated higher scores than physical workload, but along with perceived effort had the smallest range among all aspects (going from 3 to 4 across the times points). Frustration scores also had large standard deviations, and combined with the narrow range indicate that the increase in task load may not have led to any real increase in perceived frustration.

The remaining four aspects (performance, effort, mental workload, and temporal workload) all started at scores that were 4 or higher and increased to about 5.5 to 6. Perceived effort increased modestly from about 4.5 to 5.5, but unlike frustration, started at a much higher level (time point A in Figure 7) and had much smaller variability. This indicates a modest, and perhaps real, increase in perceived effort over time and task-load. Perceived temporal effort increased sharply when one variable was changed but thereafter did not exhibit any change when two variables were changed, and experienced a small reduction when all variables were changed. However, this last time point was taken after a repeat of CTN111, and may be a result of the participant gaining more practice.

Perceived performance and mental workload had the largest absolute increase in scores (from about 4 to about 6), with perceived performance increasing steadily across all time points. Perceived mental workload increased rapidly from zero to one to two variables, and thereafter either stayed the same or showed a small decline. The greatest changes in all aspects, except performance and effort, was observed when going from zero to two variables. However, any changes in perceived performance exceeded that of perceived effort.

### 5.2. EEG

From the EEG results, we can draw some conclusions about the tested groupings (which were designed to assess cognitive load) as follows:

- **Least Responsive Frequency Band:**Alpha. The alpha PSD values were not being consistently attenuated or increased with the difficulty of the task (see Figure 9). This is somewhat unexpected, as Smith and Gevins (2005) reported that frontal midline EEG attenuated alpha activity proportional to increasing cognitive load during performance of an *n*-back working memory task.
- **Most responsive electrode**: FC6. Our results are similar to those reported in a recent study by Adewale and Panoutsos (2019) which reported a power increase with workload in frontal regions. In their study, the increase was highest in FC6, next to AF4 as reported here. Further, Adewale and Panoutsos (2019) found that this region was also most responsive in the mid and high beta band, the same bands as in this study (see Table 5). The corroboration with the results reported by Adewale and Panoutsos (2019) suggests that the mid- and high-beta bands responses from FC6 require further investigation in task load experiments.
- **Most responsive frequency band**: Low Beta (12.5-18 Hz), followed by higher frequency bands. The responsiveness of the highest frequency components is only present when the T variable is considered to have greater weight in the task difficulty modulation (*Hypothetical Difficulty*). This is also implicitly tested when the variables are grouped by performance, as the main effect of the Time (T) variable was highly significant (*p* = 2*e* − 18) in the ANOVA tests for the error rate.

These findings are in accordance with Berka et al. (2007) who reported that the discriminative features for their workload classifier were mostly located in frequency bins inside the beta and gamma bands. In contrast to our findings based on monopolar recordings, their study is based on bipolar derivations, in which the CzPOz electrode contained the largest number of discriminative features. In our case, these electrodes were also selected to provide greatest discrimination among the set of electrodes (Tables 2–5).

The grouping by *Temporal Flow* could be the best approach to test for fatigue, as both time and difficulty could generate fatigue in the participant. The results from the grouping by *Time (T)* on the other hand indicate that the low Beta band may be useful in predicting detecting time pressure. A previous report has suggested that increased low-Beta power is an indicator of cognitive task demand (Ray and Cole, 1985). Thus, the *T* variable could contribute greater weight to cognitive load or task difficulty than *C* or *N* variables.

In the grouping by *Hypothetical Difficulty*, significant responses were observed over a wider range of frequencies and more numbers of electrodes, than were elicited by grouping according to temporal flow (Table 3) or Time (Table 4). This group showed greater response among all difficulty-related groupings, with significant responses across a wider range of frequency bands (in particular high-frequency bands) as well as in more parieto-central electrodes such as C4 and CP1. Similar to grouping by Time (Table 4), electrode FC6 demonstrated more significant response across all frequency bands.

We also investigated potential causes of the asymmetric frontal cortical activity. This has traditionally been associated with affective valence, specifically that increased positive (negative) affect accompanies increased left (right) cortical activity (Heller, 1990). However, Davidson (1992) postulated the approaching-withdrawing behavior in social situations as giving rise to the asymmetry; Harmon-Jones and Allen (1997) suggested that trait approach motivation was related to greater left than right frontal activity. To tease apart the various causative factors, Harmon-Jones and Allen (1998) disentangled confounds of affective valence and an approach/withdraw behavior. They showed that anger, an approach related state with *negative* valence, induced greater left cortical activity. Accordingly, it was later established that anterior asymmetric activity which favors the left hemisphere is related to approach motivation irrespective of valence (Harmon-Jones et al., 2010). We calculated an objective metric, the asymmetry index, for the alpha band. This band is inversely correlated with cerebral metabolism (Cook et al., 1998) and thus, an increase in the PSD ratio of right compared to the left hemisphere reflects increased left cortical activity.

### 5.3. Galvanic Skin Response

Skin Conductance Level (SCL, tonic response) rises in anticipation of performing tasks and fluctuates in the range of seconds to minutes depending upon the psychological state, hydration, skin dryness, and autonomic arousal. Evidence has been reported of SCL concomitant with sensitivity and the general arousal systems, as well as hippocampal information processing (Boucsein, 2012). On the other hand, Skin Conductance Response (SCR, phasic response) is typically associated with short-term events induced by discrete environmental stimuli such as sight, sound, and smell, or modulated by cognitive processes. Most SCR features were positively modulated by cognitive workload, whereas some SCL features tended to encode a rather general state of arousal – with higher emotional levels at the start and end of the experiment. The time variable (T) had a more significant phasic response than tonic response, whereas the group *C+N* did not have significant phasic or tonic responses. Likewise, phasic responses for the Hypothetical difficulty group demonstrated strongly significant responses whereas tonic responses were not significant. The *# Variables* group was the only group with significant phasic and tonic responses.

### 5.4. Eye Tracking Data

Häkkänen et al. (1999) found that bus drivers with hypersomnia have significantly higher blink duration than control groups. Morris and Miller (1996) claimed that blink duration (1) increases with time on task and (2) correlates with decreased performance. On the other hand, blink duration is reported to be correlated with mental activity and effort (Andreassi, 2013; Ikehara and Crosby, 2005). We hypothesize that in our paradigm, the negative effects on blink duration induced by cognitive measures like arousal, mental activity and interest outweigh the positive effects induced by fatigue, difficulty and drowsiness.

From a neurophysiological perspective, the pupil diameter is thought to increase with cognitive workload (Brookings et al., 1996; Kahneman and Beatty, 1966) and emotional arousal (Partala and Surakka, 2003), but to decrease with drowsiness and fatigue (Morad et al., 2000). Our measurements confirm previous findings. Also, pupil diameter was negatively correlated with task performance (*r* = −0.4), hinting that it may function as an index of fatigue and drowsiness.

In our experiments, we found that fixation duration varied significantly whether or not visual search was performed. Fixation positions were more predictable in tasks where performance was high. Blink duration correlated strongly with the participant’s individual performance, and pupil dilation was positively correlated with task difficulty.

Eyetracking data were particularly useful to infer whether visual search was necessary to solve a problem (fixation duration), but blink duration also turned out to be predictive of performance. Signal to noise ratio in fixation positions could be strongly correlated to task performance. Further, a strong positive relation between subjectively measured frustration and blink rate (*r* = 0.71) hints at the relevance of blink rate to quantify mental states, an observation that has been made very recently (Yang et al., 2017). Pupil diameter on the other hand was presumably confounded by the experimental design as cognitive load, and fatigue acted oppositely to potentially level out each other. As a consequence, its results have been found of much less significance than in related work (McCuaig et al., 2010).

### 5.5. Multimodal Data

Based on our results, we conclude that EEG was the best modality to quantify cognitive load and that the low beta band activity provides the most reliable source for prediction of task performance. The accuracy of the multimodal performance prediction model is poorer than those reported in recent studies which carried out classification, rather than regression of related quantities like operator workload (Schultze-Kraft et al., 2016). However, model selection is ultimately guided by the nature of the real-world application. Our aim here was to explore approaches where widely-different physiological variables can be integrated into statistical modeling.

Although not explored here in detail, the EEG asymmetry index may offer insights into predicting anger or frustration. However, a larger sample sample size is necessary to evaluate the possibility of differentiating these emotions in approaching and withdrawing situations. Our study suggests that this may be a promising aspect to investigate in future work.

## 6. Conclusions

In this work we investigated gradually increasing cognitive load and its correlation with several physiological variables (EEG, eye-tracking, and GSR) along with subjective data from the NASA TLX workload questionnaire. We aimed to identify pertinent information relating to cognitive load, emotional quantities like frustration, and task performance, as a function of task difficulty.

Low-beta frequency EEG waves (12.5-18 Hz) showed up more prominently as cognitive task load increased. The most responsive regions of the surface scalp EEG were found to be frontal and parietal regions. More frequent eye blinks and higher pupil dilation were detected as tasks got more difficult, while blink duration correlated strongly with task performance. Phasic components of the GSR signal were related to cognitive workload, whereas tonic components may encode a general state of arousal. Subjective data (NASA TLX) showed an increase in frustration and mental workload as reported by the participants. Based on one-way ANOVA, EEG alone, and EEG with GSR, provided the most reliable correlation to the subjective workload level and were the most informative measures for performance prediction.

Future investigations should carefully consider amending the experimental design to disentangle confounds of time and difficulty increasing simultaneously (e.g. by block randomization), thus allowing clearer inference on the dynamics of features driven by otherwise opposing forces such as pupil diameter.

## Acknowledgment

This study was supported by the research grant for the Human Centered Cyber-physical Systems Programme at the Advanced Digital Sciences Center from Singapore’s Agency for Science, Technology and Research (A*STAR). J.B. thanks the German Academic Exchange Service (DAAD) for financial support. The authors thank all the participants in this study.

## 7. Appendix

Features used for multimodal analysis:

**GSR features:** 1: SCR mean frequency, 2: SCR number, 3: SCR mean amplitude, 4: SCR sum of amplitude, 5: SCR max. amplitude, 6: SCR median, 7: SCR sum of integral (total sum of AUC), 8: Latency of first SCR after stimulus, 9: SCR maximum, 10: SCR amplitude, 11: SCL median slope, 12: SCR onset (median), 13: SCL onset (mean), 14: SCL and SCR (mean), 15: SCL amplitude, 16: SCR mean integral (mean AUC).
**Eyetracking features:** 1: Blink rate, 2: Fixation rate, 3: Saccade rate, 4: Blink duration, 5: Fixation duration, 6: Saccade duration, 7: Pupil diameter, 8: Interpupil distance, 9: Eyegaze distance, 10: POR distance, 11: Distance to screen.
**EEG features:** FC1, FC6, CP1, C4, Cz, F4, POz; each for the frequency bands: low beta, mid beta, high beta, low gamma and gamma.

